# Isoform-specific Disruption of the *TP73* Gene Reveals a Critical Role for TAp73gamma in Tumorigenesis via Leptin

**DOI:** 10.1101/2022.08.07.503085

**Authors:** Xiangmudong Kong, Wensheng Yan, Wenqiang Sun, Yanhong Zhang, Hee Jung Yang, Mingyi Chen, Hongwu Chen, Ralph W. de Vere White, Jin Zhang, Xinbin Chen

**Affiliations:** Comparative Oncology Laboratory, Schools of Veterinary Medicine and Medicine, University of California at Davis, Davis, CA, 95616; Berkeley Regional Lab, Pathology/Lab-Histology Department, The Permanente Medical group, Berkeley, CA, 94085; Pharmacology Team, Drug Discovery Center, LG Life Science R&D, LG Chem Ltd, Seoul, Korea. 150-721; Department of Pathology, University of Texas Southwestern Medical Center, Dallas, TX, 75390; Department of Biochemistry and Molecular Medicine, University of California at Davis, Davis, CA, 95616; Department of Urology Surgery, School of Medicine, University of California at Davis, Davis, CA, 95616

**Keywords:** TAp73, p53, p73 C-terminal isoforms, TAp73gamma, Leptin, oncogenesis, Obesity

## Abstract

TP73, a member of the p53 family, is expressed as TAp73 and ΔNp73 along with multiple C-terminal isoforms (α−η). ΔNp73 is primarily expressed in neuronal cells and necessary for neuronal development. Interestingly, while TAp73α is a tumor suppressor and predominantly expressed in normal cells, TAp73 is found to be frequently altered in human cancers, suggesting a role of TAp73 C-terminal isoforms in tumorigenesis. To test this, the TCGA SpliceSeq database was searched and showed that exon 11 (E11) exclusion occurs frequently in several human cancers. We also found that p73α to p73γ isoform switch resulting from E11 skipping occurs frequently in human prostate cancers and dog lymphomas. To determine whether p73α to p73γ isoform switch plays a role in tumorigenesis, CRISPR technology was used to generate multiple cancer cell lines and a mouse model in that *Trp73* E11 is deleted. Surprisingly, we found that in E11-deificient cells, p73γ becomes the predominant isoform and exerts oncogenic activities by promoting cell proliferation and migration. In line with this, E11-deficient mice were more prone to obesity and B-cell lymphomas, indicating a unique role of p73γ in lipid metabolism and tumorigenesis. Additionally, we found that *E11-*deficient mice phenocopies *Trp73*-deficient mice with short lifespan, infertility, and chronic inflammation. Mechanistically, Mechanistically, we showed that Leptin, a pleiotropic adipocytokine involved in energy metabolism and oncogenesis, was highly induced by p73γ, necessary for p73γ-mediated oncogenic activity, and associated with p73α to γ isoform switch in human prostate cancer and dog lymphoma. Finally, we showed that E11-knockout promoted, whereas knockdown of p73γ or Leptin suppressed, xenograft growth in mice. Our study indicates that the p73γ-Leptin pathway promotes tumorigenesis and alters lipid metabolism, which may be targeted for cancer management.

## Introduction

The *TP73* gene encodes a member of the p53 family of transcription factors involved in tumor suppression and development. *TP73* gene is mapped to a region on chromosome 1p36 that is frequently deleted in neuroblastoma and other tumors, thus linking its role to cancer (1). p73 shares a high similarity with p53 in the transactivation, tetramerization, and DNA binding domains, with most considerable homology between their DNA binding domains (2). Consequently, p73 activates a number of putative p53 target genes involving in cell cycle arrest, apoptosis, and differentiation (2, 3). Despite these similarities, p73 is not functionally redundant to p53. *TP73* is rarely mutated but frequently found to be up-regulated in human cancers, indicating that p73 is not a classic Knudson-type tumor suppressor (4–6). Indeed, the biological functions of p73 have been linked to neurological development, tumorigenesis, ciliogenesis, fertility, and metabolism as illustrated by *in vitro* and *in vivo* models (7–15).

The role of p73 in cancer is complicated due to the presence of multiple p73 isoforms that coordinate in a complex way to exert both oncogenic and non-oncogenic activities (15–19). Through the usage of two promoters, *TP73* encodes two classes of isoforms, TAp73 and N-terminally truncated ΔNp73. TAp73 isoforms, driven by the P1 promoter located upstream of exon 1, contain a transactivation domain similar to that in p53 and thus are suggested to function as a tumor suppressor. ΔNp73 isoforms, arising from the P2 promoter located in intron 3, lack the N-terminal transactivation domain and thus, are thought to have oncogenic potential by acting as dominant-negative inhibitors of p53 and TAp73. Mice deficient in ΔNp73 isoforms are not prone to tumors, but have defects in neurological development similar to that in total p73-KO mice (10, 11). By contrast, mice deficient in TAp73 isoforms develop spontaneous and carcinogens-induced tumors (12), suggesting a role of TAp73 as tumor suppressor. However, several recent studies also showed that TAp73 exhibits both pro-tumorigenic activities depend on the context (15, 20, 21), suggesting that the role of TAp73 in tumorigenesis is far more complex, likely being affected by the various C-terminal TAp73 isoforms.

At the C-terminus, multiple p73 isoforms are generated due to alternative splicing from exon 11 to 13 (22, 23). At transcript levels, six splicing variants (α, β, γ, δ, ε, ζ) can be detected. However, at protein levels, only three isoforms (α, β, and γ) can be detected with α being the most abundantly expressed. Due to lack of proper model systems and reagents, most studies about p73 C-terminal isoforms have been done in cell culture and thus, may not reflect the authentic biological role of these isoforms. When over-expressed, p73β showed very strong, whereas p73α and p73γ exhibited much weaker, activity to induce programmed cell death (24, 25). These differential activities are suggested due to the variation of C-terminal sequence among p73 C-terminal isoforms (23). p73α, the longest isoform, contains a sterile alpha motif (SAM) and an extreme C-terminal domain, both of which are thought to inhibit its transcriptional activity through interaction with other factors (26). Similarly, p73γ contains a unique C-terminal domain with unknown function. By contrast, p73β does not contain an extended C-terminal domain present in both p73α and p73γ. Nevertheless, these hypotheses have never been tested *in vivo*. The difficulties to study the physiological roles of p73 C-terminal isoforms are mainly due to the following reasons :1) the presence of TA/ΔN isoforms with opposing functions, which adds complications to study the C-terminal isoforms; 2) p73α is the major isoform expressed in most cells and tissues and may shield activities by other isoforms; 3) lack of specific antibodies to detect β or γ isoforms. These limitations may have led to two opposing functions for TAp73 as a tumor suppressor and a tumor promoter (27).

Leptin is a 167-amino-acid peptide hormone encoded by the Ob gene and produced by a multitude of tissues, especially adipose tissue (28). Leptin communicates with its receptor (LepR) in the hypothalamus and controls energy expenditure and appetite through a negative feedback loop, thereby maintaining the relative constancy of adipose tissue mass from being too thin or too obese (29, 30). The anti-obesity effect of Leptin was demonstrated in individuals bearing congenital leptin deficiencies (31). It was later found out that leptin therapy is ineffective as most obese individuals have a high level of endogenous plasma leptin (hyperleptinaemia) and do not respond to leptin therapy, a condition called leptin resistance (32). As a pleiotropic cytokine, leptin plays a critical role in lipid/glucose homeostasis, immune responses, hematopoiesis, angiogenesis, reproduction, and mental processes (33–36). Leptin signaling is also found to promote cancer progression, including cell proliferation, metastasis, angiogenesis and chemoresistance (37, 38). Notably, both leptin and LepR are frequently altered in many obesity-associated cancers, including lung, breast, ovarian, pancreas, liver and prostate (39–41). Thus, targeting Leptin-LepR signaling may represent a novel strategy for cancer therapeutics.

To understand the role of various p73 C-terminal isoforms in cancer development, we searched the TCGA SpliceSeq database and found that *Trp73* E11 skipping, which led to isoform switch from p73α to p73γ, frequently occurs in a subset of human cancers and dog lymphomas. Thus, to explore the biological function of p73γ isoform, we employed CRISPR technology to manipulate p73 splicing by deleting E11 in the *TP73* gene in multiple cancer cell lines and mice. We showed that E11-deficiency resulted in p73γ become the predominant isoform in both human and mouse cells. Unexpectedly, we found that p73γ, when expressed at a physiologically relevant level, did not behave as a tumor suppressor, but promoted cell migration and tumorigenesis in vitro and in vivo. Mechanistically, we showed that Leptin, a pleiotropic adipocytokine, was induced by p73γ and contributes to p73γ-mediated tumorigenesis. Moreover, we found that p73α to γ isoform switch was detected in a subset of dog lymphomas and human prostate carcinomas along with elevated expression of Leptin. Finally, we showed that targeting p73γ or Leptin led to growth suppression of E11-KO xenografts in a mouse xenograft model, suggesting that the p73γ-Leptin pathway may be explored for cancer management.

## Materials and Methods

### Reagents

Anti-Actin (sc-47778, 1:3000), anti-p130 (sc-374521, 1:3000), anti-p21 (sc-53870, 1:3000), anti-p130 (sc-374521, 1:3000) and anti-PML (sc-377390, 1:3000) were purchased from Santa Cruz Biotechnology. Anti-TAp73 (A300-126A, 1:2000) was purchased from Bethyl Laboratories. Anti-HA (MMS-101P, 1:3000) was purchased from Covance. Anti-Ki-67 (Cat#12202, 1:100), anti-B220 (Cat#70265, 1:100) and Anti-Leptin (Cat#16227, 1:500) were purchased from Cell signaling. Anti-p73γ/ε antibody was generated by Cocalico Biologicals using a peptide (PRDAQQPWPRSASQRRDEC). To-Pro-3 was purchased form Invitrogen. The WesternBright ECL HRP substrate (Cat# K12043-D20) was purchased from Advansta. Matrigel was obtained from Corning Inc. (Cat# 354230). IHC kit (Cat# PK6100) and DAB (Cat# SK4100) were purchased from Vector laboratories. Dual-Luciferase® Reporter Assay kit was purchased from Promega (Cat# E1910). Scrambled siRNA (5’- GGC CGA UUG UCA AAU AAU U -3’), p73α/γ siRNA#1 (5’- AGC CUC GUC AGU UUU UUA A -3’), p73α/γ siRNA#2 (5’- AAC CUG ACC AUU GAG GAC CUG GG -3’), Leptin siRNA#1 (5’- GCU GGA AGC ACA UGU UUA U -3’), and Leptin siRNA#2 (5’- CCA GAA ACG UGA UCC AAA UUU -3’) were purchased from Dharmacon (Chicago, IL). For siRNA transfection, RNAiMax (Life Technologies) was used according to the user’s manual. Proteinase inhibitor cocktail was purchased from Sigma-Aldrich. Magnetic Protein A/G beads were purchased from MedChem. RiboLock RNase Inhibitor and Revert Aid First Strand cDNA Synthesis Kit were purchased from Thermo Fisher. Recombinant Human Leptin Protein was purchased form R&D (398-LP).

### Plasmids

To generate a vector expressing a single guide RNA (sgRNA) that targets total p73, TAp73, ΔNp73 and E11, two 25-nt oligos were annealed and then cloned into the pSpCas9(BB) sgRNA expression vector via BbsI site (42). To generate the pSpCas9(BB)-ΔNp73-sgRNA-Blast vector expressing both blasticidin and ΔNp73 sgRNA, the blasticidin gene was amplified from pcDNA6 vector and then cloned into pSpCas9(BB)-ΔNp73 via NotI. To generate a luciferase reporter that contains the WT Leptin promoter, a DNA fragment was amplified from genomic DNA from H1299 cells, cloned into pGL2-basic vector via XmaI and HindIII sites, followed by sequence confirmation. To generate a luciferase reporter that contains mutant Leptin promoter, two-step PCR was used. The first step was to amplify two DNA fragments (Fragment #1 and #2) by using the WT Lep promoter as the template. Fragment #1 was amplified using WT forward 1 primer and mutant reverse mutant 1 primer. Fragment #2 was amplified using mutant forward 1 primer and WT reverse 1 primer. The second round of PCR was performed by using both fragment #1 and #2 as templates with a WT forward 1 and a reverse 1 primer. The resulted DNA fragment was then cloned into pGL2-basic vector via XmaI and HindIII sites. To generate a pcDNA4 vector expressing HA-tagged TAp73γ, a 680 bp DNA fragment was amplified by using cDNAs from H1299 cells as a template, and then used to replace the C-terminal of HA-TAp73α (24) via E*co*RI and X*ho*I sites. The sequences of all primers were listed in Supplemental Table S4.

### Mice and MEF isolation

*Trp73^+/−^* mice were generated as described previously (43). *E11^+/−^* mice were generated by the Mouse Biology Program at University of California at Davis. All animals and use protocols were approved by the University of California at Davis Institutional Animal Care and Use Committee. To generate WT and *E11^-/-^* MEFs, *E11^+/−^*mice were bred with *E11^+/−^* and MEFs were isolated from 12.5 to 13.5 postcoitum (p.c.) mouse embryos as described previously (44). MEFs were cultured in DMEM supplemented with 10% FBS (Life Science Technology), 55 μM β-mercaptoethanol, and 1× non-essential amino acids (NEAA) solution (Cellgro). All the genotyping primers were listed in Supplemental Table S5.

### Cell culture, Cell Line Generation

H1299 and Mia-PaCa2 cells and their derivatives were cultured in DMEM (Dulbecco’s modified Eagle’s medium, Invitrogen) supplemented with 10% fetal bovine serum (Hyclone). MCF-10A cells were cultured in DMEM:F12 (1:1) supplemented with 5% donor horse serum, 20 ng/ml EGF, 10 μg/ml insulin, 0.5 μg/ml hydrocortisone, and 100 ng/ml cholera toxin. To generate Total p73-, TAp73, ΔNP73, and E11-KO cell lines by CRISPR-Cas9, H1299 or Mia-PaCa2 cells were transfected with pSpCas9(BB)-2A-Puro vector expressing a guide RNA, and then selected with puromycin for 2-3 weeks. To generate ΔN/E11-DKO cells, E11-KO H1299 and Mia-PaCa2 cells were transfected with pSpCas9(BB)-ΔNp73-sgRNA-Blast vector, followed by selection with blasticidin for 2-3 weeks. Individual clone was picked confirmed by sequence analysis or western blot analysis. The primers used for sequencing total p73, TAp73, ΔNp73 and p73 exon 11 are listed in Supplemental Table S5. H1299 cells that inducibly express HA-tagged TAp73γ were generated as previously described (45). Briefly, pcDNA4-HA-TAp73 vector was transfected into H1299 cells in which a tetracycline repressor is expressed by pcDNA6 (46). Individual clone was picked and screened and for TAp73g expression by performing western blot analysis. To induce HA-tagged TAp73γ expression, tetracycline was added to the medium.

### Western Blot Analysis

Western blot analysis was performed as previously described (47). Briefly, whole cell lysates were harvested by 2×SDS sample buffer. Proteins were separated in 7-13% SDS-polyacrylamide gel, transferred to a nitrocellulose membrane, probed with indicated antibodies, followed by detection with enhanced chemiluminescence and visualized by VisionWorks®LS software (Analytik Jena).

### RNA isolation and RT-PCR

Total RNA was isolated with Trizol reagent as described according to user’s manual. cDNA was synthesized with Reverse Transcriptase (Promega, San Luis Obispo, CA) and used for RT-PCR. The PCR program used for amplification was (i) 94°C for 5 minutes, (ii) 94°C for 45 seconds, (iii) 58°C for 45 seconds, (iv) 72°C for 30 seconds, and (v) 72°C for 10 minutes. From steps 2 to 4, the cycle was repeated 22 times for actin and GAPDH, 28-35 times depending on the targets. All primers were used for RT-PCR are listed in Supplemental Table S6.

### ChIP Assay

ChIP assay was performed as previously described (48). Briefly, chromatin was cross-linked in 1% formaldehyde in phosphate-buffered saline (PBS). Chromatin lysates were sonicated to yield 200- to 1,000-bp DNA fragments and immunoprecipitated with a control IgG or an antibody against HA or p73γ. After reverse cross-linking and phenol-chloroform extraction, DNA fragments were purified, followed by PCR to visualize the enriched DNA fragments. The primers used for the ChIP assays were listed Supplemental Table S6.

### Luciferase reporter assay

Dual luciferase assay was performed according to the manufacturer’s instructions (Promega). Briefly, H1299 cells were plated at 5×10^4^ cells in triplicate per well in a 24-well plate and allowed to recover overnight. Cells were then transfected with the following plasmids: (1) 3 ng of pRL-SV40-Renilla; (2) 0.25 ug of a luciferase reporter, and (3) 0.25 ug of empty pcDNA3 vector or a pcDNA3 vector expressing p73α, β, or γ. The relative fold change of luciferase activity is a product of the luciferase activity induced by p73α, β, or γ divided by that induced by control vector.

### Colony formation assay

For Colony formation assay, cells (600 per well) in a six-well plate were cultured for 15 days. The cell clones were fixed with methanol/glacial acetic acid (7:1) and then stained with 0.1% of crystal violet.

### Wound healing assay

2×10^5^ cells were seeded in a 6-well plate cells and grown for 24 h. The monolayers were wounded by scraping with a P200 micropipette tip and washed two times with PBS. At indicated time points after scraping, cell monolayers were photographed with phase contrast microscopy. Cell migration was determined by visual assessment of cells migrating into the wound. Wound closure percentage was quantified using Image J plugin, Wound Healing Sizing Tool (49), by comparing the width of the wound between 0 h and indicated time points.

### Senescence assay

The senescence assay was performed as described previously (50). Briefly, primary MEFs at passage 5 were seeded at 5×10^4^ in a well of 6-well plate for 24 hours. Cells were then washed with 1×phosphate-buffered saline and fixed with 2% formaldehyde, 0.2% glutaraldehyde for 15 mins at room temperature, and then stained with fresh SA-*β*-galactosidase staining solution [1 mg/mL 5-bromo-4-chloro-3-indolyl-β-d-galactopyranoside, 40mm citric acid/sodium phosphate (pH 6.0), 5mm potassium ferrocyanide, 5 mm potassium ferricyanide, 150mm NaCl, and 2mm MgCl2]. The percentage of senescent cells was calculated as SA-β-gal positive cell divided by the total number of cells counted.

### Histological Analysis and Immunohistochemistry (IHC)

Mouse tissues were fixed in 10% (wt/vol) neutral buffered formalin, processed, and embedded in paraffin blocks. Tissues blocks were sectioned (6 μm) and stained with hematoxylin and eosin (H&E). Immunohistochemistry (IHC) analysis was performed using the Vectastain ABC Elite Kit (Vector Lab-oratories) according to manufacturer’s instruction. Briefly, tissue sections (5μm) were dewaxed and antigen-retrieved in a citrate buffer (pH 6.0), followed by incubation with a primary antibody Anti-Ki-67 (Cat#12202, 1:100), anti-B220 (Cat#70265, 1:100) or Anti-Leptin (Cat#16227, 1:500) overnight at 4 °C and then a secondary antibody for 1 h at room temperature. The slides were visualized by treatment with 3,3′-diaminobenzidine tetrahydrochloride (DAB), and then counterstained with Mayer’s hematoxylin.

### ELISA

Leptin Elisa was performed using the BioVendor Mouse and Rat Leptin Elisa kit (Cat# RD291001200R). Briefly, mouse serum was incubated with a microplate precoated with mouse leptin antibody. Bound leptin was detected with biotin labelled polyclonal anti-mouse leptin antibody conjugated to horseradish peroxidase and quantified by a chromogenic substrate at 450 nm. A standard curve is constructed by plotting absorbance values versus leptin concentrations of standards, and concentrations of unknown samples are determined using this standard curve. To measure the level of cholesterol and triglycerides, Cholesterol/Cholesterol Ester-Glo™ Assay kit (Cat#J3190, Promega) and Triglyceride-Glo™ Assay kit (Cat#J3160, Promega) from Promega were used according to user’s manual. Briefly, mouse sera or cell lysates were first incubated with cholesterol/triglycerides lysis solution at 37°C for 30 min, then incubated with cholesterol/triglycerides detection reagent for 1 hour, followed by luminescence detection with Luminometer (SpectraMAX). A standard curve is constructed and concentrations of unknown samples are determined using the standard curve.

### Three-dimensional culture for acini

The assay was performed as previously described (51). Briefly, single cell suspensions were plated onto Matrigel-coated chamber slides at 5000 cells/well in complete growth medium with 2% Matrigel and allowed to grow for 1–22 days. Overlay medium containing 2% Matrigel was renewed every 4 days. At the end of culture, cells were fixed and the nuclei were stained with 5 μg/ml of To-Pro-3 in PBS for 15 min at room temperature. Confocal microscopic images of the acinus structures were captured by the *Z*-stacking function for serial confocal sectioning at 2-μm intervals (LSM-510 Carl Zeiss laser scanning microscope) and then analyzed by Carl Zeiss software.

### Three-dimensional tumor spheroid culture

Single cell suspensions (3,000 cells/well) were plated around the rim of the well of a 96-well plate in a 4:3 mixture of Matrigel and MammoCult medium (BD Bioscience CB-40324). The cell mixture was then incubated at 37°C with 5% CO_2_ for 15 min to solidify the gel, followed by addition of 100 µL of pre-warmed MammCult. At the end of 3-D culture, cells were released from the Matrigel by incubating with 50 μL of dispase (5 mg/mL) (Life Technologies #17105-041) at 37°C for 45 min. Spheroids were imaged by a phase contrast microscope and cell viability was measured by CellTiter-Glo according to manufacturer’s guidelines (Promega).

### Transwell Migration Assay

Transwell migration assay was performed as previously described (52). Briefly, 1 x 10^5^ cells were seeded in 100 μl of serum free medium in the upper chamber of a 24-well transwell and then incubated at 37°C for 10 minutes to allow cells to settle down. Next, 600 μl of the DMEM with 10% FBS were added to the lower chamber in a 24-well transwell and cells were cultured at 37°C for various time. At the end of each time point, the transwell insert was removed from the plate and cells that had not migrated were removed by a cotton-tipped applicator. Cells that migrated to the other side of membrane were fixed with 70% ethanol for 20 minutes, stained with 0.1% of crystal violet, and photographed by phase contrast microscope. For siRNA knockdown experiment, isogenic control and E11-KO H1299 cells were transfected with a scrambled siRNA or a siRNA against p73α/γ or Leptin for 48 hours, and then subjected to transwell assays. For Leptin treatment, recombinant Leptin protein (100 ng/ml) was added to media for ten hours.

### Tissue collection

The human normal and prostate cancer specimens were obtained from Dr. Ralph De Vere White’s group, with consent from patients who underwent radical prostatectomie. Fresh frozen dog lymph nodes from clinical samples were provided by the University of California at Davis Small Animal Clinic with the owner’s permission. Samples were homogenized in Trizol, followed by RNA and protein purification according to the users’ manual.

### Xenograft assay

6 × 10^6^ cells were mixed with Matrigel (1:1 ratio) in 100 ul and then injected subcutaneously into 8-week-old BALB/c athymic nude mice (Charles River). When tumors were palpable, tumor growth was monitored for every 2 days for a period of up to 17 days. To knock down p73γ and Leptin *in vivo*, Accell siRNAs against p73γ or Leptin were synthesized from Dharmacon, which were then transiently transfected into E11-KO H1299 cells along with a scrambled Accell siRNA at a concentration of 7.5 μM. Two days post transfection, cells were collected and injected into athymic nude mice, followed by monitoring tumor growth as describe above. Mice bearing xenograft tumors also received one more intratumoral injection of Accell siRNAs (7.5 μM) at day 9. Tumor volume was calculated according to the standard formula: *V* = length × width × depth × 0.5236 (53). At the endpoint, all animals were sacrificed and the tumors weighed. One half of the tumor was stored at −80°C and the other half will be fixed in formalin and embedded with paraffin. Tumors were sectioned and H&E-stained for histopathology examination. All animals and use protocols were approved by the University of California at Davis Institutional Animal Care and Use Committee.

### Statistical analysis

The Log-rank test was used for Kaplan–Meier survival analysis. Fisher’s exact test or two-tailed students’ t test was performed for the statistical analysis as indicated. for the statistical analysis of all ELISA. Values of p< 0.05 were considered significant.

## Results

### p73α-γ switch is detected in a subset of human prostate carcinomas and dog lymphomas

Early studies have shown that p73 C-terminal isoforms are dysregulated in breast cancer and leukemia (17, 54). However, it is not clear how p73 C-terminal isoforms are altered and whether these alterations play a role in tumorigenesis. In this regard, we examined the splicing pattern of *TP73* in normal and tumor tissues by using the TCGA SpliceSeq database. Interestingly, we found that the percentage of exon 11 exclusion was much higher in Lung Squamous Cell Carcinomas (LUSCs) and Head-Neck Squamous Cell Carcinomas (HNSCs) when compared to their respective normal tissues (Fig. 1A-B). We would like to note that exon 11 skipping would lead to production of two p73 isoforms, p73γ that is switched from p73α and p73ε that is switched from p73β. Thus, to determine which p73 isoform is altered in cancer tissues, expression of p73α, p73γ, and p73ε transcripts were measured in 5 normal human prostates and 16 prostate carcinomas. We found that p73α was detectable in 5 normal prostates, undetectable in 12 prostate carcinomas, and unaltered or slightly elevated in 4 prostate carcinomas (Fig. 1C-D). By contrast, p73γ was highly expressed in prostate carcinomas as compared to that in normal prostates (Fig. 1C-D, compare lanes 1-5 with 6-13). Additionally, p73ε was almost undetectable at both normal and prostate cancer tissues (Data not shown). Similarly, we found that p73α was expressed in 3 normal dog lymph nodes but undetectable in 16 dog lymphomas (Fig. 1E, p73α panel, compare lanes 1-3 with 4-16). By contrast, p73γ expression was very low in 3 normal dog lymph nodes but elevated in all 16 dog lymphomas (Fig. 1E, p73γ panel, compare lanes 1-3 with 4-16). These data suggest that p73α isoform in normal tissues is switched to p73γ isoform in cancer tissues and that p73γ is associated with tumorigenesis.

**Figure 1.**
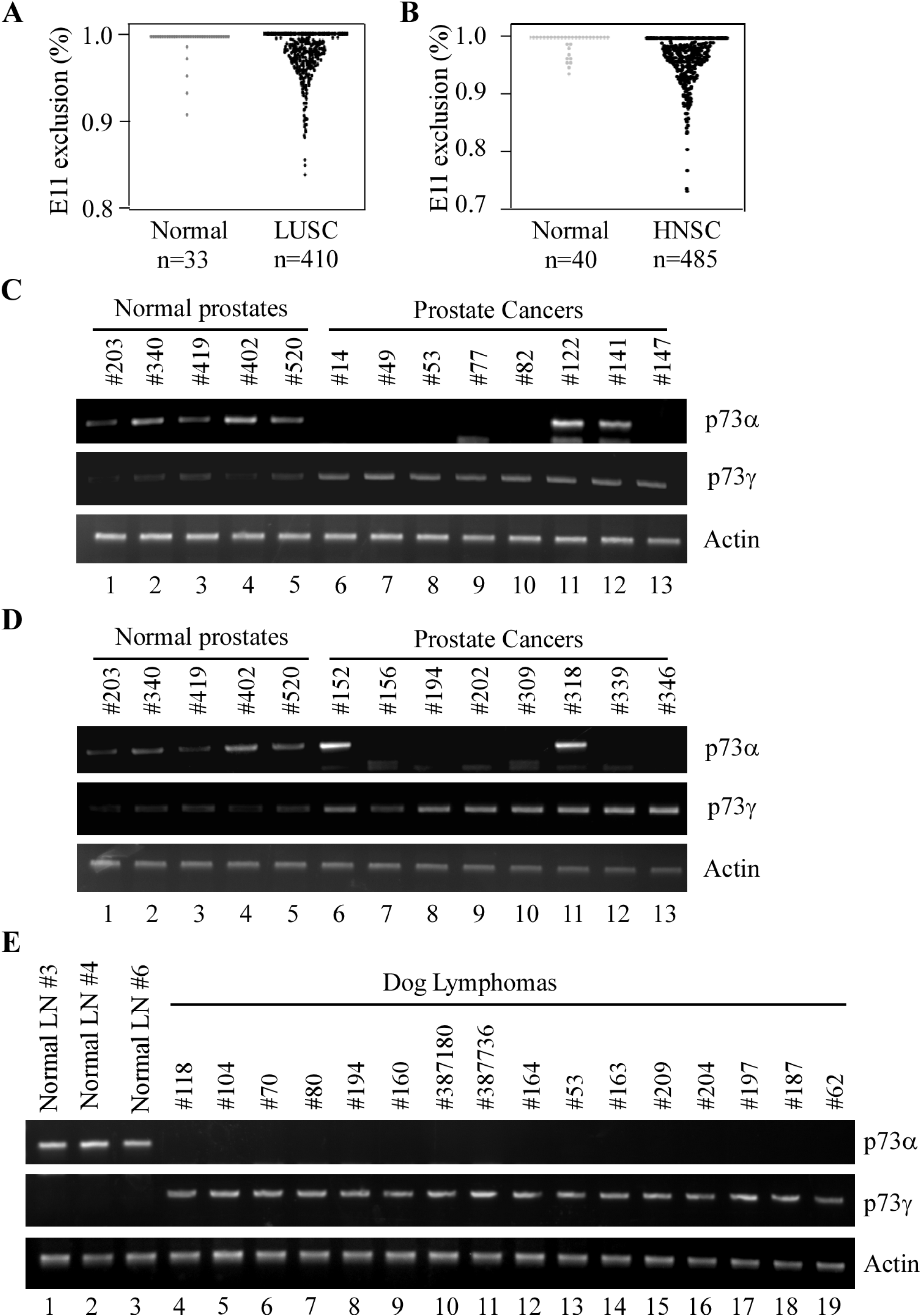
E11-skipping is detected in a subset of human cancers and dog lymphomas. **(A-B)** Percentage of *TP73* exon 11 exclusion were examined in LUSC (Lung Squamous Cell Carcinoma) and HNSC (Head-Neck Squamous Cell Carcinoma) along with their normal tissues by using TCGA SpliceSeq database. Value of 1 indicates 100% of the transcripts contains all exons, e.g p73α. Values less than 100% represents the extent of expression of the isoforms with exon 11 exclusion, e.g. *p73*γ. **(C)** The Level of p73α, p73γ, and actin transcript was examined in three normal dog lymph nodes and 16 dog lymphomas.

### Deletion of Exon 11 in the *TP73* gene leads to isoform switch from α to γ, resulting in enhanced cell proliferation and migration as well as altered epithelial morphogenesis

To mimic E11 skipping in human cancer tissue and understand the biological functions of p73γ, CRISPR-Cas9 technology was used to generate stable H1299 and Mia-PaCa2 cell lines in that the splicing acceptor for *TP73* Exon 11 (E11) was deleted by using two guide RNAs (supplemental Fig. 1A). Sequence analysis verified that E11-KO H1299 and Mia-PaCa2 cells contained a deletion of 21-nt in intron 10 and 26-nt in E11 in the *TP73* gene. We predicted that deletion of E11 splicing acceptor would trigger E11 skipping and subsequently, result in C-terminal isoform switching (Fig. 2A). Indeed, we found that when compared to isogenic control cells, E11-KO cells expressed no α/β isoforms, increased levels of γ/ ε isoforms, and unaltered δ/ζ isoforms (Fig. 2B). Additionally, TA/ΔNp73 isoforms were expressed at similar levels in both isogenic control and E11-KO H1299 cells (Fig. 2B, TA/ΔNp73 panels). The isoform switch in E11-KO cells was also confirmed with specific primers that amplify α/γ or β/ε isoforms (Supplemental Fig. 1B-C).

**Figure 2.**
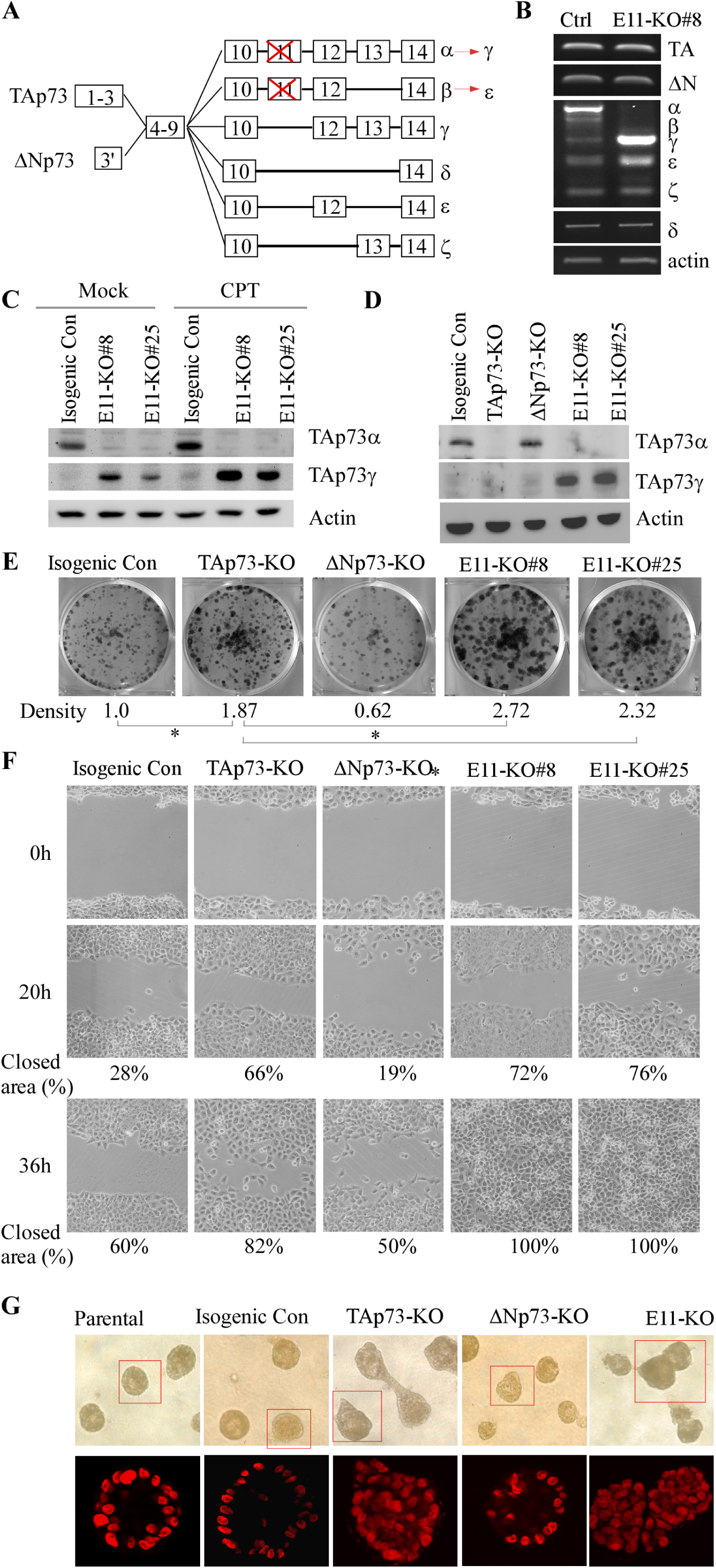
Deletion of Exon 11 in the *TP73* gene leads to isoform switch from α to γ, resulting in enhanced cell proliferation and migration as well as altered epithelial morphogenesis. **(A)** Schematic representation of p73 isoforms and isoform switch resulting from E11 deletion. **(B)** The level of p73 splicing variants was examined by RT-PCR in isogenic control and E11-KO H1299 cells. **(C)** The level of TAp73α, TAp73γ, and actin protein was measured in isogenic control and E11-KO H1299 cells mock-treated or treated with camptothecin (CPT). **(D)** The level of TAp73α, TAp73γ, and actin protein was measured in isogenic control, TAp73-KO, ΔNp73-KO, and E11-KO H1299 cells. **(E)** Colony formation was performed with isogenic control, TAp73-KO, ΔNp73-KO, and E11-KO H1299 cells. The was quantified as relative density. **(F)** Scratch assay was performed with isogenic control, TAp73-KO, ΔNp73-KO, and E11-KO H1299 cells. Phase contrast photomicrographs were taken immediately after scratch (0 h), 20h, and 36h later to monitor cell migration. **(G)** Representative images of parental, isogenic control, TAp73-KO, DNp73-KO, and E11-KO MCF10A cells in three-dimensional culture. Top panel: phase contrast image of acini in 3-D culture. Bottom panel: Confocal images of cross-sections through the middle of acini stained with TO-PRO-3.

Due to lack of specific antibodies and relative low abundance, p73γ is almost undetectable under normal conditions. Thus, a peptide (PRDAQQPWPRSASQRRDE) derived from the unique C-terminal region in p73γ/ε was used to generate an antibody, called anti-p73γ/ε antibody, which was found to recognize p73γ/ε but not p73α (Supplemental Fig. 1D). We showed that in isogenic control cells, TAp73α was the mostly abundant protein expressed as detected by an anti-TAp73 antibody, whereas in E11-KO cells, TAp73γ became the predominantly expressed isoform and was detected by anti-p73γ/ε antibody under normal and DNA-damage induced conditions (Fig. 2C and Supplemental Fig. 1E). Nevertheless, other p73 isoforms, including p73β and p73ε, remained undetectable likely due to their low abundance. These data confirmed that deletion of the acceptor site for E11 in *TP73* led to E11 skipping and subsequently, resulted in isoform switch from p73α in isogenic control cells to p73γ in E11-KO cells.

By using two different promoters, p73 C-terminal isoforms are expressed as TA/ΔNp73 proteins, which are known to have opposing functions (17). Thus, when studying the p73 C-terminal isoforms, it is also important to consider the N-terminal variations. To this end, stable TAp73- or ΔNp73-KO H1299 and Mia-PaCa2 cell were generated (Fig. 2D and Supplemental Fig. 1F) and used as controls to study the function of E11 deficiency. We found that knockout of TAp73 enhanced, whereas knockout of ΔNp73 reduced, colony formation as compared to isogenic control cells (Fig. 2E), consistent with previous reports (55, 56). Interestingly, the number of colonies formed by E11-KO cells was higher than that by TAp73-KO cells, becoming the highest among all four cell lines (Fig. 2E). To verify this, tumor sphere formation assay was performed and showed that loss of TAp73 or E11 promoted, whereas loss of ΔNp73 inhibited, cell proliferation in H1299 cells (Supplemental Fig. 1G). Moreover, to examine whether E11 deficiency affects cell migration, scratch assay was performed. We found that knockout of TAp73 promoted, whereas knockout of ΔNp73 slightly inhibited, cell migration in H1299 and Mia-PaCa2 cells in a time-dependent manner (Fig. 2F and Supplemental Fig. 1H), consistent with previous observations (56, 57). Notably, E11-KO cells migrated even faster than isogenic control or TAp73-KO H1299 and Mia-PaCa2 cells (Fig. 2F and supplemental Fig. 1H). Consistently, transwell migration assay showed that E11-KO H1299 cells exhibited a markedly enhanced ability to transmigrate through a membrane when compared to isogenic control, TAp73-KO, and ΔNp73-KO H1299 cells (Supplemental Fig. 1I). Furthermore, to determine whether E11 deficiency has an effect on epithelial morphogenesis, E11-KO, TAp73-KO and ΔNp73-KO MCF10A cells were generated (Supplemental Fig. 1J) and then subjected to three-dimensional (3-D) culture. We found that parental and isogenic control MCF10A cells underwent normal cell morphogenesis (acini with hollow lumen) (Fig. 2G, left two panels) whereas *ΔNp73*-KO showed delayed acinus formation (small acini) (Fig. 2G, ΔNp73 panel). Interestingly, E11-KO and TAp73-KO showed aberrant cell morphogenesis (irregular acini with filled lumen) (Fig. 2G, *TAp73^-/-^* and E11-KO panels). These data suggest that loss of E11 promotes oncogenesis by altering epithelial cell morphogenesis and enhancing cell proliferation and migration.

### TAp73γ is primarily responsible for the oncogenic effects observed in E11-KO cells

The enhanced potential of cell proliferation and migration in E11-KO cells could be mediated by increased TAp73γ expression or simply by loss of TAp73α as similar phenotypes were also observed in TAp73-KO cells albeit to a less extent (Fig. 2E-G and Supplemental Fig. 1G-I). Thus, to characterize the role of TAp73γ in oncogenesis, stable H1299 cell lines in that TAp73γ can be inducibly expressed under the control of a tetracycline-inducible promoter were generated (Supplemental Fig. 2A). We showed that upon induction, TAp73γ promoted cell migration (Supplemental Fig. 2B), consistent with the data from E11-KO cells (Fig. 2F and Supplemental Fig. 1G-I). Next, to determine the role of endogenous p73γ in cell proliferation and migration, two p73α/γ siRNAs (si-p73α/γ#1 or #2), which were designed to target the junction region of E12-E13 or E13-E14 in p73α/γ mRNA (Supplemental Fig. 3A), were synthesized and found to efficiently knock down p73α and p73γ in isogenic control and E11-KO cells (Fig. 3A-B). Next, cell migration was measured by Scratch and Transwell assays. We showed that knockdown of p73α/γ enhanced cell migration in isogenic control cells (Fig. 3C and Supplemental Fig. 3B). By contrast, knockdown of p73α/γ decreased cell migration in E11-KO cells (Fig. 3C and Supplemental Fig. 3B). Similarly, we found that knockdown of p73α/γ enhanced cell viability of tumor spheres in isogenic control cells but decreased in E11-KO cells (Supplemental Fig. 3C). These data suggested that cell proliferation and migration is inhibited by p73α, the major isoform in isogenic control cells, but enhanced by p73γ, the major isoform in E11-KO cells.

**Figure 3.**
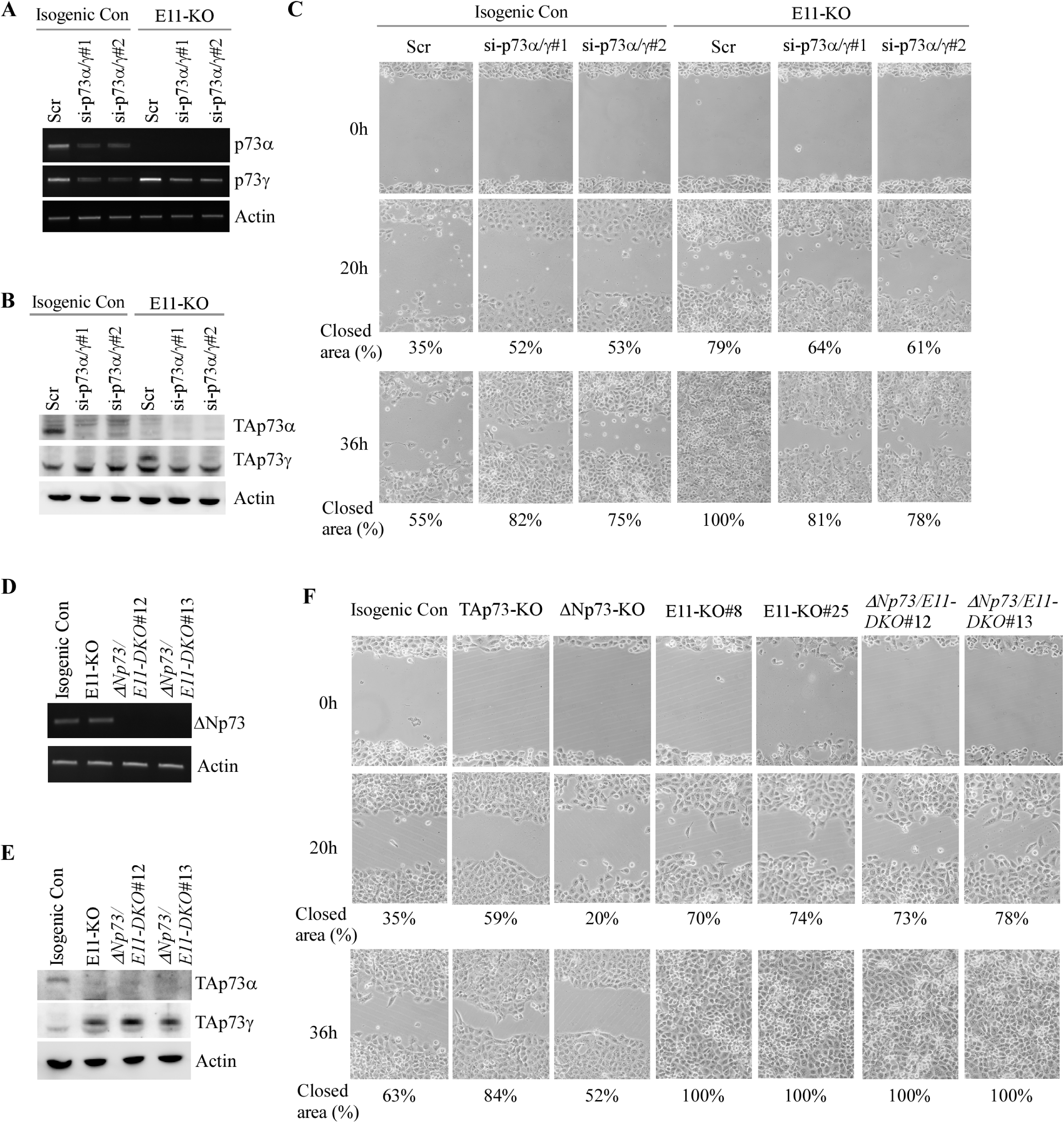
TAp73γ is primarily responsible for the oncogenic effects observed in E11-KO cells. **(A)** Isogenic control and E11-KO H1299 cells transfected with a scrambled siRNA or siRNAs against p73α/γ for 3 days, followed by RT-PCR analysis to measure level of p73α, p73γ, and actin transcripts. **(B)** Cell were treated as in (A) and then subjected to western blot analysis to measure level of TAp73α, TAp73γ, and actin proteins using antibodies against TAp73, p73α/γ and actin. **(C)** Scratch assay was performed with isogenic control and E11-KO H1299 cells transfected with a scrambled siRNA or siRNAs against p73α/γ for 3 days. Phase contrast photomicrographs were taken immediately after scratch (0 h), 20h, and 36h later to monitor cell migration. **(D)** The Level of ΔNp73 and actin transcripts was measured in isogenic control, E11-KO, ΔNp73/E11-DKO H1299 cells. **(E)** The Level of TAp73α, TAp73γ, and actin proteins was measured by western blot analysis using isogenic control, E11-KO, ΔNp73/E11-DKO H1299 cells. **(F)** Scratch assay was performed with isogenic control, TAp73-KO, ΔNp73-KO, E11-KO, and ΔNp73/E11-DKO H1299 cells. Phase contrast photomicrographs were taken immediately after scratch (0 h), 20h, and 36h later to monitor cell migration.

p73γ can be expressed as two isoforms, TAp73γ and ΔNp73γ. To determine which p73γ isoform is responsible for the enhanced cell migration in E11-KO cells, we generated a dual-KO (DKO) cell line by deleting all ΔNp73 isoforms in E11-KO H1299 and Mia-PaCa2 cells. We showed that ΔNp73 transcript was not expressed, whereas a similar amount of TAp73γ protein was expressed, in ΔNp73/E11-DKO H1299 and Mia-PaCa2 cells as compared to that in E11-KO cells (Fig. 3D-E and Supplemental Fig. 3D-E). We found that the ability of cells to migrate was not altered by ΔNp73/E11-DKO as compared to that by E11-KO (Fig. 3F and Supplemental Fig. 3F). Considering that knockout of ΔNp73 inhibited cell proliferation and migration (Fig. 2E-F and Supplemental Fig. 1G-H), we conclude that TAp73γ was able to overcome the inhibitory effect of ΔNp73 deficiency on cell proliferation and migration. Together, these data suggest that TAp73γ is primarily responsible for the enhanced cell proliferation and migration observed in E11-KO cells.

### Deletion of E11 in the *Trp73* gene leads to shortened lifespan, increased incidence of spontaneous tumor and chronic inflammation in mice

CRISPR-Cas9 was used to generate a mouse model in that E11 in the *Trp73* gene was deleted (Supplemental Fig. 4A). A cohort of WT, *E11^+/-^*, and *E11^-/-^* MEFs was then generated (Supplemental Fig. 4B) and used to examine whether E11 deficiency would lead to isoform switch in mice. We found that although MEFs mainly expressed p73α/γ/ζ transcripts, E11 deficiency led to isoform switch from p73α in WT MEFs to p73γ in both *E11^+/-^* and *E11^-/-^* MEFs (Fig. 4A), which was similar to that in human cells (Fig. 2B). Next, SA-β-gal staining was performed to evaluate whether E11 deficiency had an effect on cellular senescence, an intrinsic mechanism of tumor suppression. Surprisingly, *E11^-/-^*MEFs were less prone to cellular senescence as compared to WT MEFs (Fig. 4B), along with decreased expression of senescence markers, including PML, p130, and p21 (Fig. 4C).

**Figure 4.**
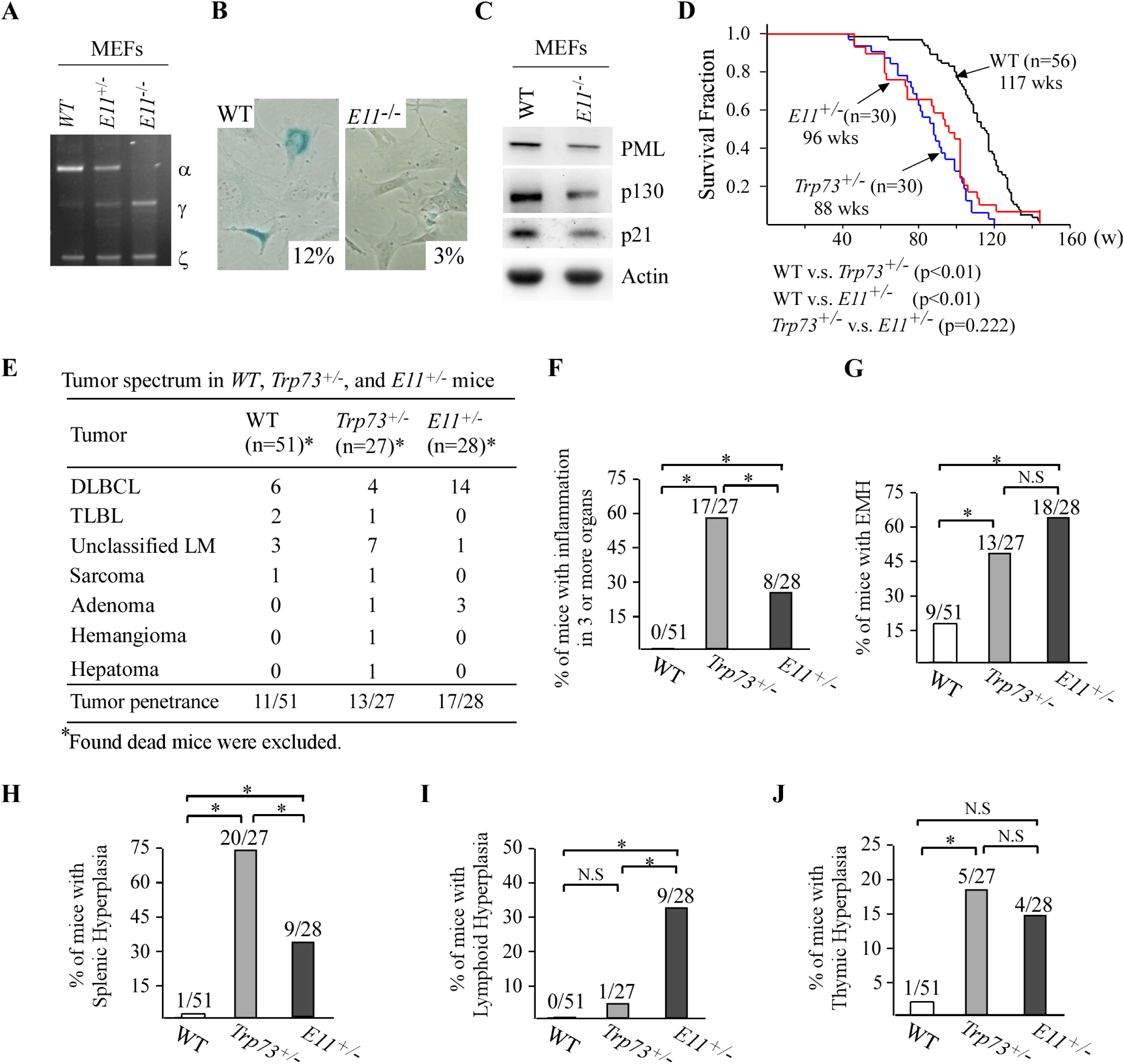
Deletion of E11 in the *Trp73* gene leads to shortened lifespan, increased incidence of spontaneous tumor and chronic inflammation in mice. **(A)** The level of Trp73α, Trp73γ, Trp73ζ and actin transcripts were measured in WT, *E11^+/-^*, and *E11^-/-^* MEFs by RT-PCR analysis. **(B)** SA-β-gal staining was performed with WT and *E11^-/-^* MEFs. The percentage of SA-β-gal-positive cells was shown in each panel. **(C)** Western blot was performed to measure the level of PML, p130, p21, and actin proteins in WT and *E11^-/-^* MEFs using antibodies against PML, p130, p21, and actin. **(D)** Kaplan-Meyer survival curves of WT (n=56), *Trp73^+/-^* (n=30), and *E11^+/-^* (n=30) mice. **(E)** Tumor spectra in WT (n=56), *Trp73^+/-^* (n=27), and *E11^+/-^* (n=28) mice. **(F)** Percentage of WT, *Trp73^+/-^*, and *E11^+/-^* mice with inflammation in 3 or more organs. **(G)** Percentage of WT, *Trp73^+/-^*, and *E11^+/-^* mice with extramedullary hematopoiesis. **(H)** Percentage of WT, *Trp73^+/-^*, and *E11^+/-^* mice with splenic hyperplasia. **(I)** Percentage of WT, *Trp73^+/-^*, and *E11^+/-^* mice with lymphoid follicular hyperplasia. **(J)** Percentage of WT, *Trp73^+/-^*, and *E11^+/-^* mice with thymic hyperplasia.

To examine the biological functions of E11 deficiency *in vivo*, a cohort of *E11^+/-^* and *E11^-/-^* mice were generated. We found that *E11^-/-^*mice were runty, infertile and prone to hydrocephalus (Supplemental Fig. 4C and data not shown). Thus, *E11^-/-^* mice exhibited similar phenotypes as *Trp73^-/-^* mice (7, 14) and were not suitable for long-term tumor study. However, young *E11^+/-^* mice appeared to be normal. In this regard, a cohort of *E11^+/-^*mice was generated and monitored for potential abnormalities along with WT and *Trp73^+/-^* mice. We would like to mention that to minimize the number of animals used, all WT and 26 out of 30 *Trp73^+/-^* mice, which were generated previously but had same genetic background and maintained in the same facility (43, 58, 59), were used as controls. We found that the lifespan for *E11^+/-^*mice (96 weeks) and *Trp73^+/-^* mice (88 weeks) was much shorter than that for WT mice (117 weeks) (Fig. 4D). However, there was no difference in lifespan between *Trp73^+/-^* and *E11^+/-^*mice (Fig. 4D). Pathological analysis indicated that 11 out of 51 WT mice, 13 out of 27 *Trp73^+/-^* mice, and 17 out of 28 *E11^+/-^* mice developed spontaneous tumors (Fig. 4E; Supplemental Tables 1-3). Statistical analysis indicated that the tumor incidence was significantly higher for both *Trp73^+/-^* and *E11^+/-^* mice as compared to that for WT mice (p=0.0211 for WT vs. *Trp73^+/-^*; p=0.0011 for WT vs. *E11^+/-^* by Fisher’s exact test). Interestingly, although there was no difference in tumor incidence between *E11^+/-^* and *Trp73^+/-^*mice, the percentage of diffuse large B cell lymphoma (DLBCL) was significantly higher in *E11^+/-^* mice (50%) than that in *Trp73^+/-^* mice (14.8%) (Fig. 4E and Supplemental Fig. 4D) (p=0.009 by Fisher exact test). These data suggest a role of p73γ in promoting B-cell lymphomagenesis, which is consistent with the observation that p73γ is frequently up-regulated in dog lymphomas (Fig. 1E).

In addition to tumors, *E11^+/-^* mice developed other pathological defects, including chronic inflammation, EMH, and hyperplasia in lymph node, thymus or spleen. Specifically, 17 out of 27 *Trp73^+/-^*and 8 out of 28 *E11^+/-^* mice, whereas 0 out of 51 WT mice, showed chronic inflammation in 3 or more organs (Fig. 4F; Supplemental Fig. 4E and Tables 1-3). The chronic inflammation was much less severe in *E11^+/-^*mice as compared to *Trp73^+/-^* mice (Fig. 4F), suggesting that p73γ partially compensates p73α in suppressing inflammatory response. Moreover, when compared to WT mice, *Trp73^+/-^* and *E11^+/-^* mice were prone to EMH (Fig. 4G). Furthermore, *Trp73^+/-^* and *E11^+/-^* mice showed various degree of hyperplasia in immune organs, such as lymph node, spleen and thymus when compared to WT mice (Fig. 4H-J). Briefly, the percentage of splenic hyperplasia was higher in *Trp73^+/-^* and *E11^+/-^* mice than that in WT mice, with *Trp73^+/-^* even higher than *E11^+/-^*mice (Fig. 4H). The percentage of lymphoid hyperplasia was higher in *E11^+/-^* mice than that in WT and *Trp73^+/-^* mice (Fig. 4I), whereas the percentage of thymic hyperplasia was higher in *Trp73^+/-^*mice than that in WT and *E11^+/-^* mice (Fig. 4J). Together, these data indicated that *E11*-deficient mice phenocopies *Trp73*-deficient mice in short lifespan, infertility, chronic inflammation and tumor incidence, indicating that p73γ cannot compensate p73α for tumor suppression and fertility.

### *E11*-deficient mice are prone to obesity

We noticed that *E11^+/-^* mice were bigger and fattier as compared to WT mice. We also found that the size of adipocytes was larger in both male and female *E11^+/-^* mice than that in WT mice (Fig. 5A). Likewise, a significant increase in visceral fat (VAT) mass was observed in both male and female *E11^+/-^*mice as compared to WT mice (Fig. 5B-C). Moreover, when compared to WT mice, *E11^+/-^*but not *Trp73^+/-^* mice showed increased incidence in liver steatosis as characterized by lipid deposition in the hepatocytes (Fig. 5D-E). Consistently, the levels of transcripts for alanine aminotransferase (ALT), aspartate aminotransferase (AST), and γ-glutamyltransferase 1 (GGT1) (60, 61), all of which are associated with liver steatosis, were increased in *E11^+/-^* livers when compared to WT liver (Supplemental Fig. 5A). These observations let us speculate that E11 deficiency promotes obesity in mice. To test this, the level of serum cholesterol and triglycerides was measured and found to be much higher in *E11^+/-^* mice than that in WT and *Trp73^+/-^*mice at ∼65 weeks (Fig. 5F-G), suggesting that E11 deficiency enhances lipid storage. Consistent with this, younger *E11^+/-^* mice at 20 or 32 weeks also showed increased body weights (Fig. 5H-I) and elevated levels of serum cholesterol and triglycerides (Fig. 5J-K) when compared to WT mice. Furthermore, to examine whether p73γ is responsible for decreased lipid catabolism observed in *E11*^+*/-*^ mice, the level of cholesterol and triglycerides was measured in isogenic control and E11-KO cells with or without knockdown of p73γ. We found that E11 deficiency led to marked increase in the level of cholesterol and triglycerides in both H1299 and Mia-PaCa2 cells (Fig. 5L-N and Supplemental Fig. 5B-D). However, upon knockdown of p73γ, elevated level of cholesterol and triglycerides by E11-KO was reduced to normal level in isogenic control cells (Fig. 5L-N and Supplemental Fig. 5B-D). By contrast, the level of cholesterol and triglycerides was increased by knockdown of p73α/γ in isogenic control H1299 and Mia-PaCa2 cells (Fig. 5L-N and Supplemental Fig. 5B-D), suggesting a role of p73α in inhibiting lipid storage. Together, these data suggest that *E11* deficiency leads to obesity in mice, which is largely induced by TAp73γ.

**Figure 5.**
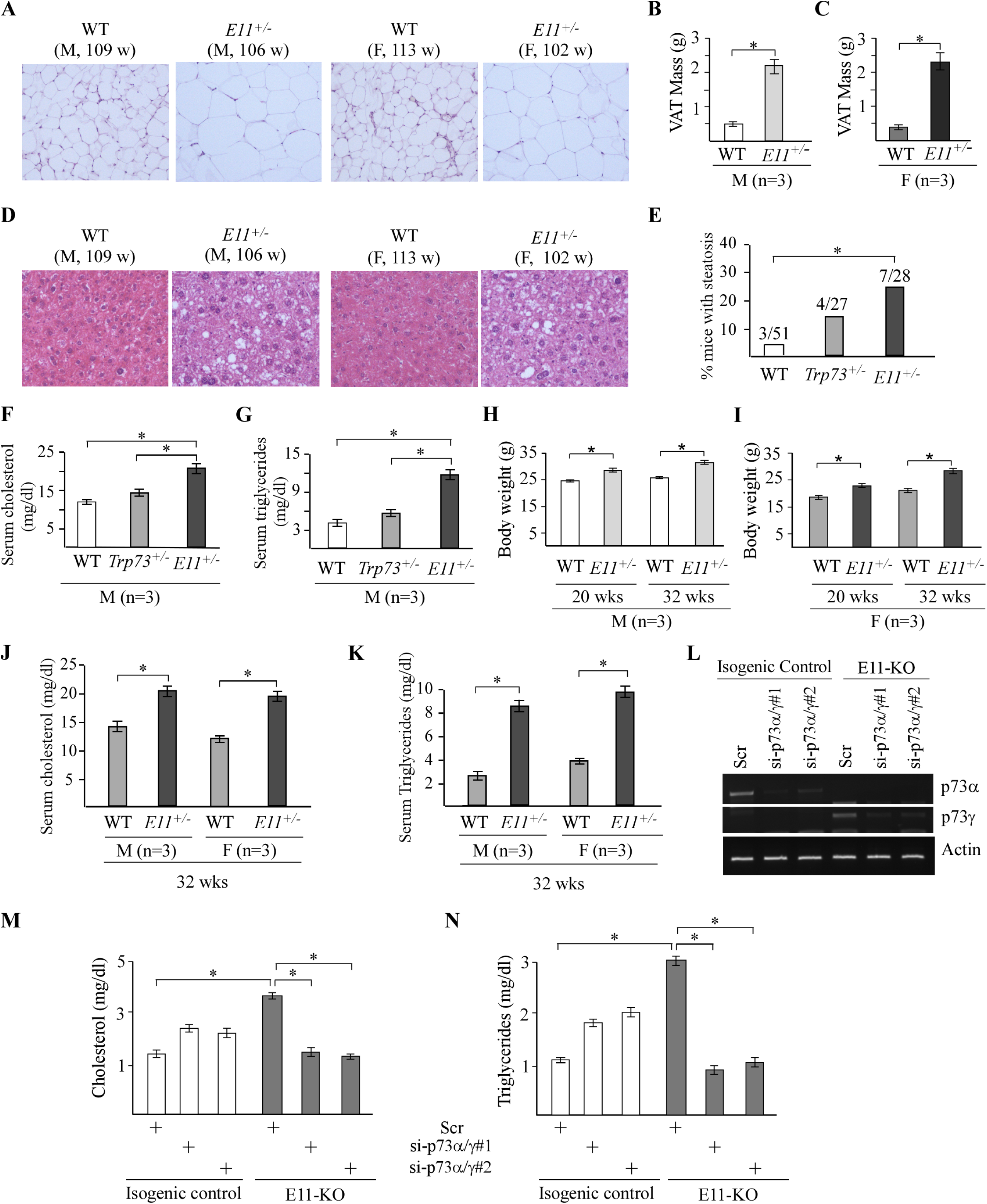
E11-deficient mice are prone to obesity. **(A)** Representative images of H.E stained visceral adipose tissues from male and female WT and *E11^+/-^* mice. **(B-C)** VAT mass of male (B) and female (C) WT and *E11^+/-^* mice at ∼100 weeks. **(D)** Representative images of H.E stained liver tissues from male and female WT and *E11^+/-^* mice. **(E)** Percentage of WT, *Trp73^+/-^*, and *E11^+/-^* mice with liver steatosis. **(F-G)** The level of serum cholesterol (F) and triglycerides (G) was measured in male WT, *Trp73^+/-^*, and *E11^+/-^* mice at 100 weeks. **(H-I)** Body weights were measured in male (H) and female (I) WT, *Trp73^+/-^*, and *E11^+/-^* mice at 20 or 32 weeks. **(J-K)** The level of serum cholesterol (J) and triglycerides (K) was measured in male and female WT, *Trp73^+/-^*, and *E11^+/-^* mice at 32 weeks. **(L)** The level of p73α, p73γ, and actin transcripts was measured in isogenic control and E11-KO H1299 cells transfected with a scrambled siRNA or siRNAs against p73α/γ. **(M-N)** Cells were treated as in (L), followed by measurement of cholesterol (M) and triglycerides (N).

### E11 deficiency leads to elevated production of Leptin, a novel target of TAp73γ

Several studies have shown that dysfunctional adipose tissues secret adipocytokines, such as leptin, and subsequently, promote adipose tissue and systemic inflammation, which is considered to be responsible for many obesity-related complications, including cancer (62–64). Interesting, we found that *E11^+/-^* but not WT mice showed pronounced low-grade inflammation in visceral adipose tissues (VATs) (Supplemental Fig. 6A) as well as increased expression of several proinflammatory cytokines, such as IL6, IL1α and TNFα, in VAT and liver (Fig. 6A). These observations prompted us to speculate whether E11 deficiency leads to altered expression of leptin, a pleiotropic adipocytokine that is known to be associated with obesity, adipose tissue inflammation and cancer (65–67). In addition to adipocytes, Leptin was found to be expressed in normal epithelial and carcinoma cells and subsequently, promotes tumor growth through autocrine and paracrine signaling (68, 69). Indeed, we found that the level of leptin transcript was increased by E11 deficiency in the VAT and liver tissues as well as in MEFs (Fig. 6A and Supplemental Fig. 6B). Consistent with this, IHC assay showed an elevated expression of leptin protein in *E11^+/-^* kidney and liver as compared to WT kidney and liver, respectively (Supplemental Fig. 6C). Moreover, the level of Leptin protein in the VAT and serum was highly increased in age- and gender-matched *E11^+/-^* mice as compared to that in WT and *Trp73^+/-^*mice (Fig. 6B-C). While Leptin expression was increased in *Trp73^+/-^* mice, it was still much lower than that in *E11^+/-^* mice (Fig. 6B-C). Furthermore, the level of serum Leptin was found to be increased in both male and female *E11^+/-^* mice at very young ages of 20- or 32-week-old when compare to WT mice (Fig. 6D-E).

**Figure 6.**
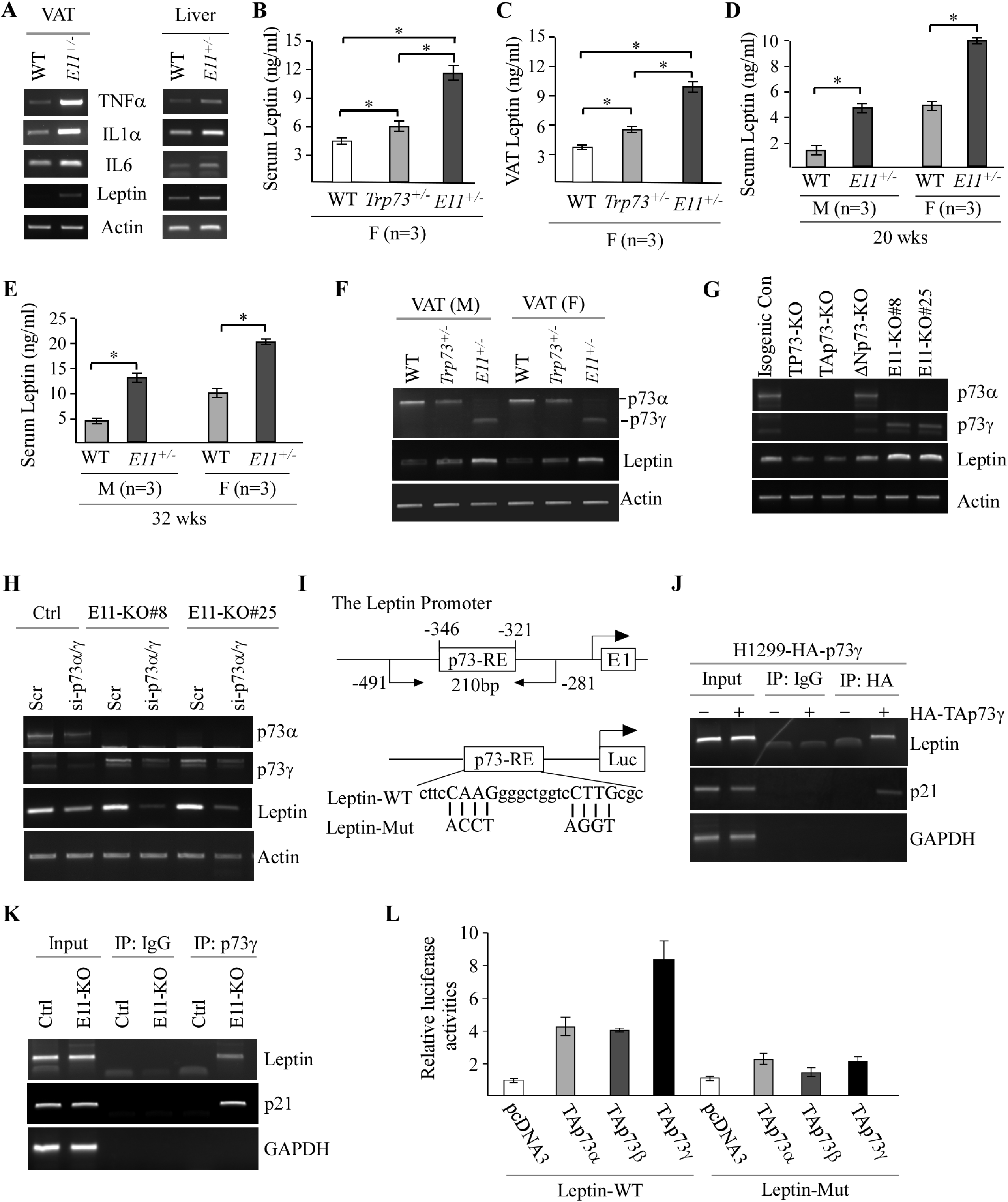
E11 deficiency leads to elevated production of Leptin, a novel target of TAp73γ. **(A)** The level of TNFα, IL1, IL-1α, Leptin, and actin transcripts was measured in VAT and liver from WT and *E11^+/-^* mice. **(B-C)** The level of serum (B) and VAT (C) Leptin was measured in female WT, *Trp73^+/-^*, and *E11^+/-^* mice at 65 weeks. **(D-E)** The level of serum leptin was measured in male and female WT and *E11^+/-^* mice at 20 (D) or 32 (E) weeks. **(F)** The level of p73, Leptin, and actin transcripts was measured in VATs from male and female WT, *Trp73^+/-^*, and *E11^+/-^* mice. **(G)** The level of p73α, p73γ, Leptin, and actin transcripts was measured isogenic control, total p73-KO, TAp73-KO, DNp73-KO, and E11-KO H1299 cells. **(H)** The levels of p73α, p73γ, Leptin, and actin transcripts was measured isogenic control and E11-KO H1299 cells transfected with a scrambled siRNA or siRNA against for p73α/γ for 3 days. **(I)** Schematic representation of the Leptin promoter, the location of p73-RE as well as luciferase reporters with WT or mutant p73RE. **(J)** H1299 cells were uninduced or induced to express TAp73γ for 24 hours, followed by ChIP assay to measure the binding of TAp73γ to the Leptin, p21 and GAPDH promoter. **(K)** Isogenic control and E11-KO cells were used for ChIP assay to examine the binding of p73γ to Leptin and p21 promoter. **(L)** Luciferase assay was performed with H1299 cells transfected with a control vector or vector expressing TAp73α, TAp73β, and TAp73γ. The relative fold-change of luciferase activity was calculated as a ratio of luciferase activity of each construct versus an empty vector.

The observations above let us speculate whether increased expression of p73γ in E11-deficient mice contributes to Leptin expression. To test this, Leptin transcript was measured in VATs from age- and gender-matched WT, *Trp73^+/-^*, and *E11^+/-^* mice. We found that the level of Leptin transcript was slightly increased in *Trp73^+/-^* mice, but markedly increased in *E11^+/-^*mice (Fig. 6F), suggesting that Leptin expression is induced by p73γ. To verify this, Leptin transcript was measured in TP73-KO, TAp73-KO, ΔNp73-KO, and E11-KO H1299 cells. We found that knockout of all p73 or TAp73 decreased, whereas knockout of ΔNp73 had little effect on, the level of Leptin transcript (Fig. 6G). However, the level of Leptin was much higher in E11-KO cells when compared to isogenic control, TP73-, TAp73-, and ΔNp73-KO H1299 cells (Fig. 6G). We also showed that ectopic expression of TAp73γ led to increased expression of Leptin (Supplemental Fig. 6D) whereas knockdown of p73γ markedly decreased the level of Leptin transcript in E11-KO H1299 and Mia-PaCa2 cells (Fig. 6H and Supplemental Fig. 6E). Interestingly, knockdown of p73α/γ also led to reduction of Leptin expression in isogenic control cells in that p73α is the predominant isoform (Fig. 6H and Supplemental Fig. 6E), suggesting that p73α can regulate Leptin transcription. To further test this, we found that ectopic expression of TAp73α, TAp73β, or TAp73γ alone was able to induce Leptin expression, with strongest induction by TAp73γ (Supplemental Fig. 6F).

To determine whether Leptin is a direct target of TAp73, we searched the Leptin promoter and found a potential p73 response element (p73-RE) located at nt -346 to -321 (Fig. 6I). Thus, ChIP assay was performed and showed that ectopic TAp73γ was able to bind to the LEP promoter (Fig. 6J). As a positive control, TAp73γ bound to the p21 promoter (Fig. 6J), a bona fide target of the p53 family (70). Consistent with this, endogenous p73γ were able to bind to the Leptin promoter in E11-KO H1299 cells as compared to isogenic control cells (Fig. 6K). Furthermore, to determine whether the p73-RE (nt -346 to -321) is required for TAp73 to transactivate the Leptin promoter, luciferase reporters that contain WT or mutant p73-RE were generated (Fig. 6I). We found that TAp73α/β/γ were able to activate the Leptin-WT reporter, with the strongest activation by TAp73γ (Fig. 6L). By contrast, all TAp73 isoforms were unable to transactivate the luciferase reporter containing mutant p73-RE (Fig. 6L). Together, these data indicate that Leptin is a novel target of TAp73γ and may play a role in TAp73γ-induced oncogenic activities.

### Leptin is a critical mediator of TAp73γ in oncogenesis and altered lipid metabolism

Leptin is known to play a critical role in oncogenesis through altered lipid metabolism (71–73). Thus, we examined whether knockdown of Leptin has any effect on the level of cholesterol and triglycerides by using two different Lep siRNAs in both H1299 and Mia-PaCa2 cells (Fig. 7A and Supplemental Fig. 7A). We showed that knockdown of Leptin abrogated the elevated level of cholesterol and triglycerides by E11-KO in both H1299 and Mia-PaCa2 cells (Fig. 7B-C and Supplemental Fig. 7B-C), consistent with the observation that knockdown of p73γ also abrogated the elevated level of cholesterol and triglycerides by E11-KO (Fig. 5M-N and Supplemental Fig. 5C-D). Next, we examined the role of Leptin in cell migration and showed that in isogenic control cells, cell migration was moderately inhibited by knockdown of Leptin in both H1299 (Fig. 7D-E and Supplemental Fig. 7D) and Mia-PaCa2 cells (Supplemental Fig. 7E-F). By contrast, the enhanced cell migration by E11-KO was abolished by knockdown of Leptin in both H1299 (Fig. 7D-E and Supplemental Fig. 7D) and Mia-PaCa2 cells (Supplemental Fig. 7E-F), consistent with the observation that knockdown of p73γ also abolished the enhanced cell migration by E11-KO (Fig. 3C and Supplemental Fig. 3B). To further analyze the role of Leptin in p73γ-mediated cell migration, we asked whether supplementation of Leptin would have an effect on cell migration. To make sure that the effect of Leptin is transmitted through its receptor, we measure the expression of Leptin receptor (LepR). We found that LepR was expressed at a similar level across isogenic control, TAp73-KO and E11-KO H1299 cells (Supplemental Fig. 7G), suggesting that these cells have an intact Leptin signaling pathway and that LepR expression is not regulated by p73. Importantly, we found that cell transmigration was enhanced by treatment with Leptin in isogenic control, TAp73-KO and E11-KO H1299 cells (Supplemental Fig. 7H). Furthermore, to determine whether the p73γ-Leptin axis plays a role in cancer progression, we examined whether the expression pattern of Leptin is correlated with that of p73γ in human prostate carcinomas and dog lymphomas. We found that Leptin expression was low in normal human prostates but highly elevated in prostate carcinomas, which correlates well with the expression of p73γ (Fig. 7F-G, Leptin and p73γ panels, compare lanes 1-5 with 6-13). Similarly, Leptin and p73γ transcripts were found to be highly expressed coordinately in dog lymphomas as compared to normal dog lymph nodes (Fig. 7H, Leptin and p73γ panels, compare lanes 1-3 with 4-19). These data suggest that Leptin is a mediator of TAp73γ in oncogenesis and altered lipid metabolism and that the TAp73γ-Leptin pathway plays a role in the development of prostate cancer and lymphoma.

**Figure 7.**
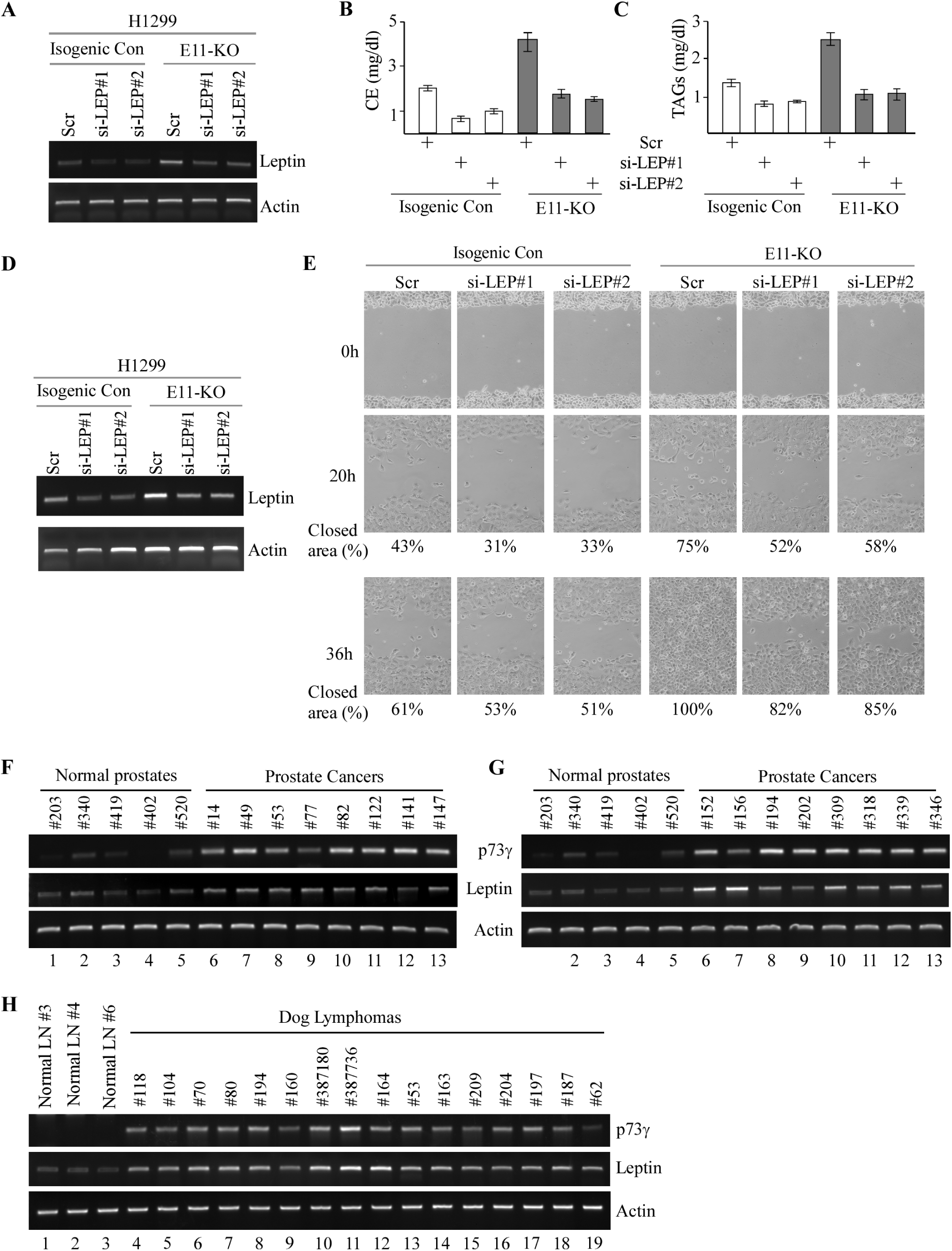
Leptin is a critical mediator of TAp73γ-mediated oncogenesis and altered lipid storage. **(A and D**) The level of Leptin and actin transcripts was measured in isogenic control and E11-KO H1299 cells transfected with a scrambled siRNA or siRNAs against Leptin. **(B-C)** The level of cholesterol (B) and triglycerides (C) was measured in cells treated as in (A). **(E)** Scratch assay was performed with H1299 cells treated as in (D). Phase contrast photomicrographs were taken immediately after scratch (0 h), 20h, and 36h later to monitor cell migration. (**F-G**) The level of p73γ, Leptin and actin transcripts was examined in five normal prostates and 16 human prostate carcinomas. **(H)** The Levels of p73γ, Leptin and actin were examined in three normal dog lymph nodes and 16 dog lymphomas.

### Targeting p73γ or Leptin inhibits tumor growth *in vivo*

To explore the therapeutic potential of targeting the p73γ-Leptin pathway for cancer management, xenograft models were established by using isogenic control and E11-KO H1299 cells. We showed that loss of E11 led to enhanced tumor growth (Fig. 8A). Consistent with this, tumor size and weight were significantly greater for E11-KO group than that for control group (Fig. 8B-C). We also showed that the levels of p73γ and Leptin transcripts and proteins were much higher in E11-KO tumors than that in WT tumors (Fig. 8D and supplemental Fig. 8A). Additionally, H.E analysis showed that both WT and E11-KO H1299 xenografts were human cells (Fig. 8E). Next, we determined whether the elevated levels of p73γ and Leptin are responsible for the enhanced tumor growth by E11-KO. To test this, siRNAs against p73γ or Leptin were synthesized along with a scrambled siRNA as a control. We found that the rate of growth for E11-KO tumors injected with p73γ or Leptin siRNA was much slower than the ones injected with the control scrambled siRNA (Fig. 8F). Consistent with this, the tumor size and weight were much less for E11-KO tumors injected with p73γ or Leptin siRNA than the ones injected with the control scrambled siRNA (Fig. 8G-H). As expected, we showed that both p73γ and Leptin were efficiently knocked down by their respective siRNAs (Fig. 8I and Supplemental Fig. 8B-C). Notably, we found that treatment with p73γ or Leptin siRNA led to extensive tumor necrosis in xenografts as compared with that treated with the control scrambled siRNA (Fig. 8J). Together, we showed that E11-KO promotes, whereas knockdown of p73γ or Leptin suppresses, xenograft growth in mice.

**Fig. 8.**
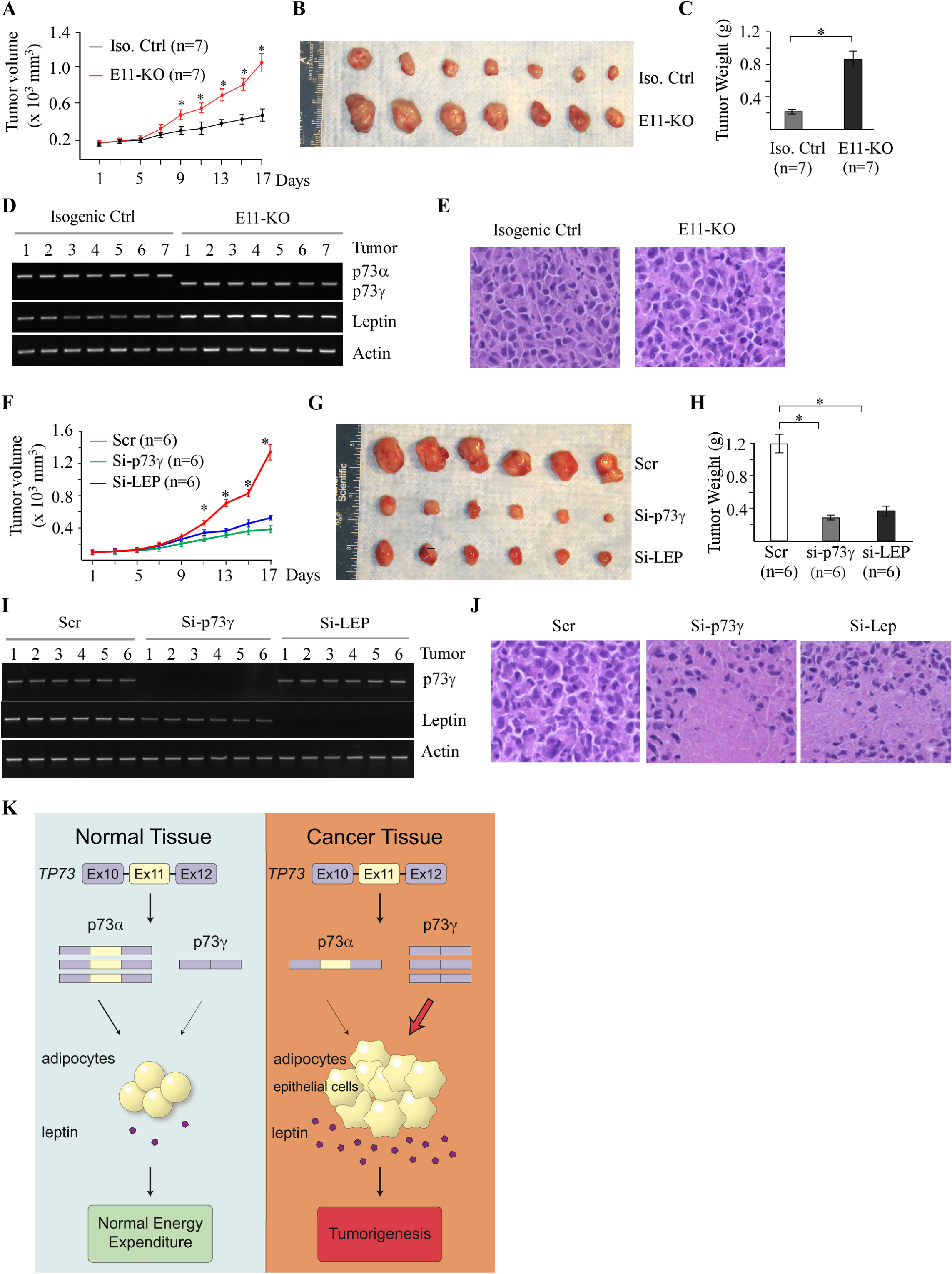
Targeting p73γ or Leptin inhibits tumor growth *in vivo*. **(A)** Xenograft tumor growth in nude mice from isogenic control or E11-KO H1299 cells (Error bars represent SEM, * indicates p<0.05 by student t test). **(B)** Images of tumors excised from isogenic control and E11-KO groups. **(C)** Tumor weigh distribution between isogenic control and E11-KO groups upon termination of tumor growth experiments at day 17. (* p<0.05 by Student’s t-test). **(D)** The levels of p73α, p73γ, Leptin, and actin transcripts were measured in the tumors from isogenic control and E11-KO groups. **(E)** Representative images of H.E-stained tumors from isogenic control and E11-KO groups. **(F)** Xenograft tumor growth in nude mice from E11-KO H1299 cells transfected with scrambled siRNA or a siRNA against Leptin or p73γ (Error bars represent SEM, * indicates p<0.05 by student t test). **(G)** Images of tumors excised from E11-KO groups with or without knockdown of Leptin or p73γ. **(H)** Tumor weigh of E11-KO tumors with or without knockdown of Leptin or p73γ (* p<0.05 by Student’s t-test). **(I)** The levels of p73α, p73γ, Leptin, and actin transcripts were measured in E11-KO tumors with or without knockdown of Leptin or p73γ. **(J)** Representative images of H.E-stained E11-KO tumors with or without knockdown of Leptin or p73γ. **(K)** A proposed model to elucidate the role of p73α/γ switch and the p73γ-Leptin axis in tumorigenesis.

## Discussion

The biology of p73 is complicated because p73 is expressed as multiple N- and C-terminal isoforms, some of which may have opposing functions. While TA and ΔNp73 isoforms are shown to exert opposing activities in tumorigenesis by multiple *in vitro* and *in vivo* model systems, the biological significance of C-terminal p73 splicing variants (α, β and γ) remains largely uncharacterized. In this study, we aimed to manipulate alternative splicing of *TP73* using CRISPR-Cas9. We showed that deletion of E11 in *TP73* leads to isoform switch from p73α to p73γ in both human cancer cells and mice. Importantly, we found that owing to p73γ expression, E11 deficiency leads to enhanced tumorigenesis and obesity *in vitro* and *in vitro*. Moreover, we identified Leptin, a key hormone in energy homeostasis, as a novel target of p73 and a critical mediator of TAp73γ in oncogenesis and aberrant lipid metabolism. Furthermore, we showed that p73α-γ switch is detected in a subset of dog lymphomas, along with elevated expression of Leptin. Finally, to explore the therapeutic potential of the p73γ-Leptin pathway, we showed that E11-KO promotes, whereas knockdown of p73γ or Leptin suppresses, H1299 xenograft growth in mice. These results have provided compelling evidence and prompted us to hypothesize that p73 C-terminal isoforms, that is, TAp73α and TAp73γ, can exert two opposing functions in tumorigenesis in a manner similar to that by TA and ΔN p73 isoforms. These results may have also explained the discrepancies in several early studies that TAp73 have both pro- and anti-tumorigenic activities (15, 19–21, 27, 74–76), which are exerted by TAp73α and TAp73γ, respectively. A model to elucidate the role of p73α/γ switch and the p73γ-Leptin axis in tumorigenesis is proposed and shown in the Fig. 8J.

We and others have shown that ectopic expression of TAp73α and TAp73γ can inhibit cell growth albeit to a much less extent when compared to p53 or TAp73β (24, 25, 77, 78), suggesting that TAp73α and TAp73γ have a role in tumor suppression. Here, we showed that when expressed at a physiological relevant level *in vivo*, TAp73γ possesses an oncogenic activity whereas TAp73α has a tumor suppressive function. Specifically, we showed that knockout or knockdown of TAp73α, the major isoform in H1299 and MiaPaCa2 cells, leads to enhanced cell proliferation and migration (Figs. 2E-G and 3C, Supplemental Fig. 1G-I). By contrast, E11 deficiency, which switches TAp73α to TAp73γ, leads to enhanced cell proliferation and migration as well as xenograft tumor growth, which can be attenuated by knockdown of p73γ (Fig. 2E-G, 3C, and 8, Supplemental Fig. 1G-I, 3B-C, and 3F). Moreover, mice deficient in *Trp73*, presumably p73α, were prone to spontaneous tumors, such as lymphoma (Fig. 4E), consistent with previous observations that loss of *Trp73* promotes Myc-induced B-cell lymphomas or mutant p53-mediated T-cell lymphomas (43, 79). By contrast, E11-deficient mice were prone to spontaneous tumors, such as DLBCLs (Fig. 4E and supplemental Table 3), presumably due to elevated p73γ expression in these mice. Furthermore, we showed that while p73α was detectable in normal tissues, p73γ was elevated in a subset of dog lymphomas (Fig. 1C). These data suggest that p73α and p73γ have opposing functions in tumorigenesis and that isoform switch from p73α to p73γ is a critical event for cancer progression. Indeed, we found that percentage of exon 11 exclusion, which would produce p73γ isoform, was much higher in tumor samples as compared to normal tissues (Fig. 1A-B). In line with this, TAp73γ transcript was found to be aberrantly expressed in human hematological malignancies, such as leukemia and non-Hodgkin’s lymphomas (80–82). Moreover, one study showed that in breast cancer, p73α and p73β are the main isoforms expressed in normal breast tissues whereas other C-terminal splice variants, including p73γ, are mainly detected in breast cancers (54). Thus, further studies are warranted to examine the expression profile of p73γ in various types of cancers. Moreover, it is important to identify the splicing factor that regulates the isoform switch from p73α to p73γ, which would further our understanding of p73 splicing variants in cancer development.

In addition to tumorigenesis, we noticed that p73α and p73γ have distinct and overlapped roles in multiple pathological processes based on the phenotypes from *Trp73*- and *E11*-deficient mice. We reasoned that if p73γ can compensate for loss of p73α, no abnormalities would be observed in *E11*-deficient mice. Instead, we found that both *Trp73*- and *E11*-deficient mice were runty, infertile, and prone to hydrocephalus (Supplemental Fig. 4C and data not shown) (7). We also found that *Trp73*-deficient mice and to a lesser degree, *E11*-deficient mice were susceptible to chronic inflammation and splenic hyperplasia (Fig. 4F and 4H). These data indicate that p73α has distinct roles in hydrocephalus, fertility, chronic inflammation and splenic hyperplasia, which cannot be fully substituted by p73γ. On the other hand, p73γ appeared to have a prominent role in lipid metabolism. Specifically, we found that *E11*-deficient mice were prone to spontaneous liver steatosis, increased body weight and VAT mass, and elevated serum triglycerides and cholesterol, all of which were not observed in *Trp73*-deficient mice (Fig. 5A-E). To verify the role of p73γ in lipogenesis, we showed that the elevated level of cholesterol and triglycerides by E11-KO was abrogated by knockdown of p73γ (Fig. 5L-N and Supplemental Fig. 5B-D). Together, these data suggest that p73α and p73γ may have opposing roles in lipid metabolism in that p73γ promotes, whereas p73α inhibits, lipogenesis.

In this study, we found that Leptin expression is induced by p73γ, leading to elevated level of Leptin in serum, VAT, and possibly many other tissues (Fig. 6A and Supplemental Fig. 6B-C). We also found that accumulation of cholesterol/triglycerides and enhanced cell migration by E11 deficiency are abrogated by knockdown of Leptin (Fig. 7B-C and 7E; Supplemental Fig. 7B-F), which phenocopies the effect of p73γ-knockdown in E11-KO cells (Fig. 3C and Supplemental Fig. 1G-I and 5C-D). In line with this, we found that Leptin treatment was able to promote cell transmigration in isogenic control, TAp73-KO and E11-KO H1299 cells (Supplemental Fig. 7H), indicating that Leptin functions as a mediator of p73 in promoting cell migration. To uncover the mechanism, we showed that Leptin is a target of p73γ (Fig. 6F-L). Interestingly, we noticed that while capable, p73α is much weaker than p73γ to induce Leptin expression (Fig. 6L and Supplemental Fig. 6F). One possibility is that the unique C-terminal domains in p73α and p73γ can recruit distinct sets of transcription coactivators and thus, differentially regulate gene expression, including Leptin, which warrants further investigation.

To explore the physiological significance of the p73γ-Leptin pathway, we found that Leptin expression correlates well with isoform switch from p73α to p73γ in a set of dog lymphomas (Fig. 7F). Leptin was initially found to control energy homeostasis through the hypothalamus and was given a high hope for treating obesity (83). However, it was found later that both too little and too much Leptin can lead to obesity (84, 85), referred to as “leptin resistance” (86, 87). At this moment, it is uncertain whether Leptin directly contributes to obesity mediated by p73γ, partially due to the above-mentioned paradoxical roles of Leptin in obesity. However, since obesity is now considered a risk factor for many cancers, the p73γ-Leptin pathway may provide a physiological link between cancer and obesity. Indeed, it should be noted that many studies, including data from this study, indicate that Leptin and its receptor are expressed in both normal and tumor tissues and that Leptin is found to promote inflammation, tumor growth, and angiogenesis (71, 88). Considering that the p73γ is highly expressed in tumors such as dog lymphoma (Fig. 1C), it is possible that p73γ plays a critical role in activating the Leptin signaling pathway in tumors, which subsequently promotes tumorigenesis. Moreover, we found that knockdown of p73γ or Leptin was able to inhibit growth of E11-KO xenograft tumors (Fig. 8), which would open a new revenue for cancer management by targeting the p73γ-Leptin pathway. Together, these data indicate that as a target of p73, Leptin mediates p73γ in tumor promotion and altered lipid metabolism and that the p73γ-Leptin pathway can be targeted for cancer management.

## Conclusion

Our studies indicate that p73α-γ switch may be a common phenomenon during cancer development and that TAp73α and TAp73γ exhibit two opposing functions in tumorigenesis. Our studies also provide compelling evidence that p73γ has an oncogenic activity and that the p73γ-Leptin pathway plays a critical role in promoting tumorigenesis and lipid metabolism, which can be explored as a novel cancer treatment strategy by blocking Leptin signaling in p73γ-expressing tumors.

## Abbreviations

E11: exon 11
SAM: Sterile alpha motif
LepR: Leptin Receptor
TCGA: The Cancer Genome Atlas
CRISPR: Clustered regularly interspaced short palindromic repeats
MEF: Mouse Embryonic Fibroblast
KO: knockout
LUSC: Lung squamous cell carcinoma
HNSC: Head-neck squamous cell carcinoma
WT: Wild type
EMH: Extramedullary hematopoiesis
ALT: Alanine aminotransferase
AST: aspartate aminotransferase
GGT1: γ-glutamyltransferase 1
VAT: Visceral adipose tissue
IL6: Interleukin 6
IL1α: Interleukin 1α
TNFα: Tumor necrosis factor α
IHC: Immunohistochemistry
p73-RE: p73 response element
DLBCL: Diffuse large b cell lymphoma

## Ethical Approval

All animals and use protocols were approved by the University of California at Davis Institutional Animal Care and Use Committee.

## Authors’ contributions

J.Z., X.C., and X.K. designed the research. X.K., W.Y., W.S., Y.Z., H.J.Y. and J.Z. performed the research. J.Z., M.C., H.C., and X.C. analyzed the data. J.Z., X.C., and X.K. wrote the paper. All authors approved the submitted and final versions of the manuscript.

## Availability of data and materials

The authors confirm that the data supporting the findings of this study are available within the article and its supplementary materials.

